# Prius: From Differentiated Genes to Affected Pathways

**DOI:** 10.1101/038901

**Authors:** Shailesh Patil, Bharath Venkatesh, Randeep Singh

**Affiliations:** SAP Labs India, 138, EPIP Zone, Whitefield, Bengaluru, Karnataka 560066

## Abstract

Expression analysis and variant calling workflows are employed to identify genes that either exhibit a differential behaviour or have a significant functional impact of mutations. This is always followed by pathway analysis which provides greater insights and simplifies explanation of observed phenotype. The current techniques used towards this purpose have some serious limitations. Only a small number of genes which satisfy certain thresholds are used for pathway analysis. All the shortlisted genes are treated as equal ignoring the differences in p-values and fold changes. These genes are treated as independent entities and interactions among them are ignored for statistical pathway analysis. Hence, there is serious disconnect between the techniques employed and networked nature of the data. Various Pathway data ases have great degree of discordance on structure of pathway graphs. Many of the pathways are still far from complete. Current algorithms do not take into account this uncertainty. In this paper, we propose a theoretical framework *Prius* to overcome many limitations of current techniques. Prius perturbs the gene expression fold changes through interaction network and generates an ordered list of affected pathways. Thus, it integrates the networked nature of the data and provides facility to weigh each gene differently and in the process we do away with the need of arbitrary cut-offs. This framework is designed to be modular and provides the researchers with flexibility to plug analytical tools of their choice for every component. We also demonstrate effectiveness of our approach for personalized and cohort analysis of cancer gene expression samples with PageRank as one of the modules in the framework. The R package for Prius is available on GitHub.

## 1 Introduction

Gene expression analysis and structural variants detection tools are used to identify genes that are significantly affected in a given disease condition. Tools ranging from earlier generation microarrays to Next Generation Sequencing like RNAseq, DNAseq and Exomeseq are often used to achieve this task. Microarray and RNAseq gene expression experiments are performed to measure changes in gene expression levels across conditions like normal vs tumor. Statistical analysis of this data usually results in p-value and fold change for each gene. Cutoffs are applied on both p-value (usually less than 0.05) and absolute fold change (usually greater than 2) to declare some of the genes as statistically significant. Structural Variant analysis workflows detect the variants in the given sample and various tools like SIFT [1], PolyPhen2 [2] and MutationAssessor [3] are employed to identify the functional impact of the mutations. Sometimes these are converted to gene level p-values and some genes are shortlisted based on certain thresholds. However, a list of significant genes alone does not provide necessary explanatory power owing to the complex interaction network of these genes. Therefore, these significant genes are further used to detect significantly affected pathways. Pathways are essentially groupings of genes based on their interactions and functions. Each pathway represents a subgraph of the gene interaction network and represents a specific cellular functionality. Techniques such as Over-Representation Analysis [4], Functional Class Scoring [5] and Pathway Topology based analysis are used to identify significantly affected pathways[6].

### 1.1 Need for a New Pathway Analysis Framework

Though a great variety of methodologies are available for pathway analysis, there are serious inconsistencies in these current approaches. Two recent papers by Khatri et al. [6] and by Mitrea et al. [7] provide a detailed review of various analytical approaches along with their shortcomings. The following are some recurring issues

1. The p-value depends on the nature of the test performed, type of multiple testing correction and more importantly degrees of freedom.
2. Reliable tests are not available for personalized analysis using a normal tumor sample pair.
3. The cutoffs are rigid and therefore a lot of information is lost. For example, a gene of p-value 0.05 and fold change 2 qualifies to be statistically significant whereas gene with p-value 0.051 and fold change 4 is not considered for further downstream analysis.
4. All the significantly expressed genes are treated equally for their pathways analysis irrespective of differences in their fold change values.
5. Pathways analysis treats and tests each pathway independently. However pathways interact with each other and many genes are part of more than one pathway.
6. Many of the current pathway databases do not concur on the graph structure of the pathways and the interactions graphs are far from complete. This uncertainty is not modeled by current algorithms

To summarize, the current methodologies use ad-hoc cutoffs, use only part of the information and do not model the interactive nature of the data.

## 2 A Novel Approach

To address the problems mentioned in the previous section, we propose a new approach ***Prius*** which accommodates information of every single gene in the experiment and models the network of interactions of the genes. Following are three abstract components of the model.

1. A measure of gene’s affectedness which is treated as disturbance. This could be fold change for expression experiments or functional impact of mutations.
2. A mechanism for perturbation of the disturbance through the interaction network. This mechanism needs to converge in finite number of steps and provide a ranking of genes.
3. A mechanism to calculate pathway’s deviation in terms of rank changes of constituent genes. This mechanism is expected to order pathways by deviations.

Prius uses the PageRank algorithm [8] as perturbation mechanism. PageRank has a proven mathematical foundation [9] and has been successfully applied to variety of network analysis problems across domains including biology [10].

The model is very general and each component can be modeled in a variety of different ways depending on nature of the task at hand. We illustrate this with some examples for each component in section 4.

We first define the notation that we use in our paper, followed by the description of PageRank. We then discuss the model - the components and their interpretation.

### 2.1 Notation

Table 1 shows the notation that we follow in this paper.

**Table I.**
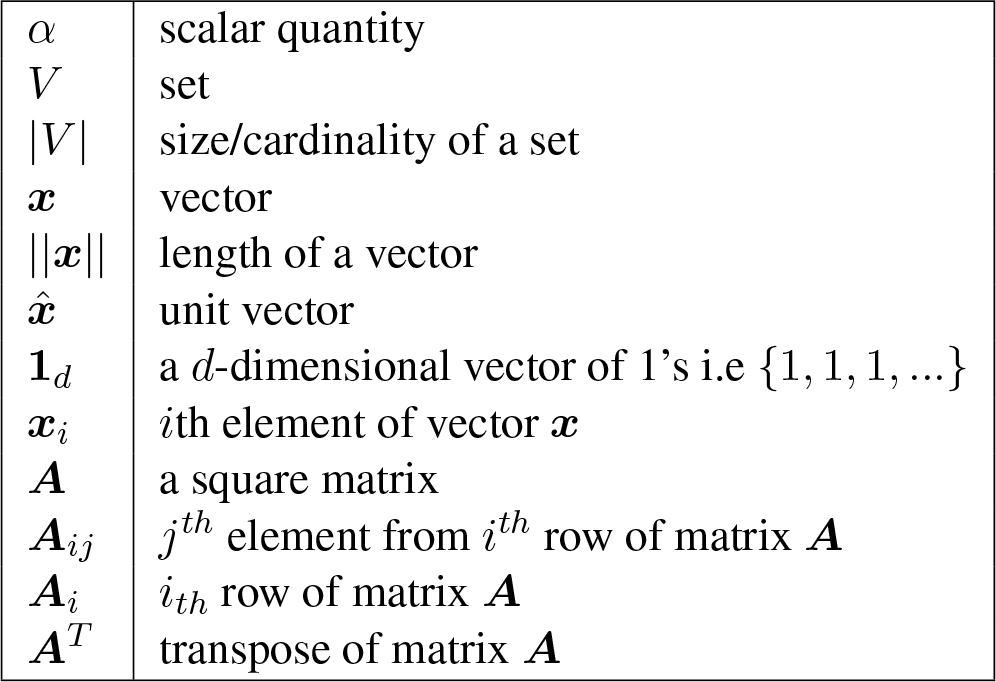
Notation

Small letters are used to denote scalar quantities. Uppercase letters are used to denote sets. We use bold type faces to denote matrices and vectors. Bold uppercase letters are used to represent a matrix while bold lowercase letters are used to represent vectors. Subscripts are used to denote individual components.

### 2.2 PageRank

Given a directed, weighted graph *G*(*V, E*) consisting of the set of edges *E* which represent the interactions between vertices (genes) in the set *V*, the PageRank vector measures the importance of each vertex.

Each element of the adjacency matrix of the graph *G* is given by -

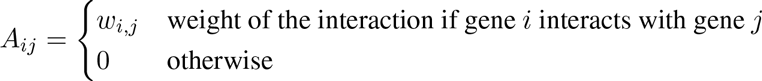

If the weights of the interactions are not available, the matrix *A_ij_* reduces to a boolean matrix with *w_ij_* = 1 for each interaction.

The PageRank vector is computed on the normalized adjacency matrix *M* of the graph, where each entry is divided by the sum of its row (also called the outdegree of the vertex). The properties of *M* are -

1. 0 ≤ *M_ij_* ≤ 1
2. *M_ij_* = 0 if gene *i* does not interact with gene *j*
3. 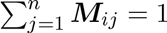

The | V |-dimensional PageRank vector *r* is computed iteratively as the solution to following equation

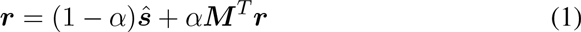

PageRank and personalized PageRank are illustrated using a toy 11-vertex network in Figure 1. The vertices are sized according to their PageRank

**Figure 1.**
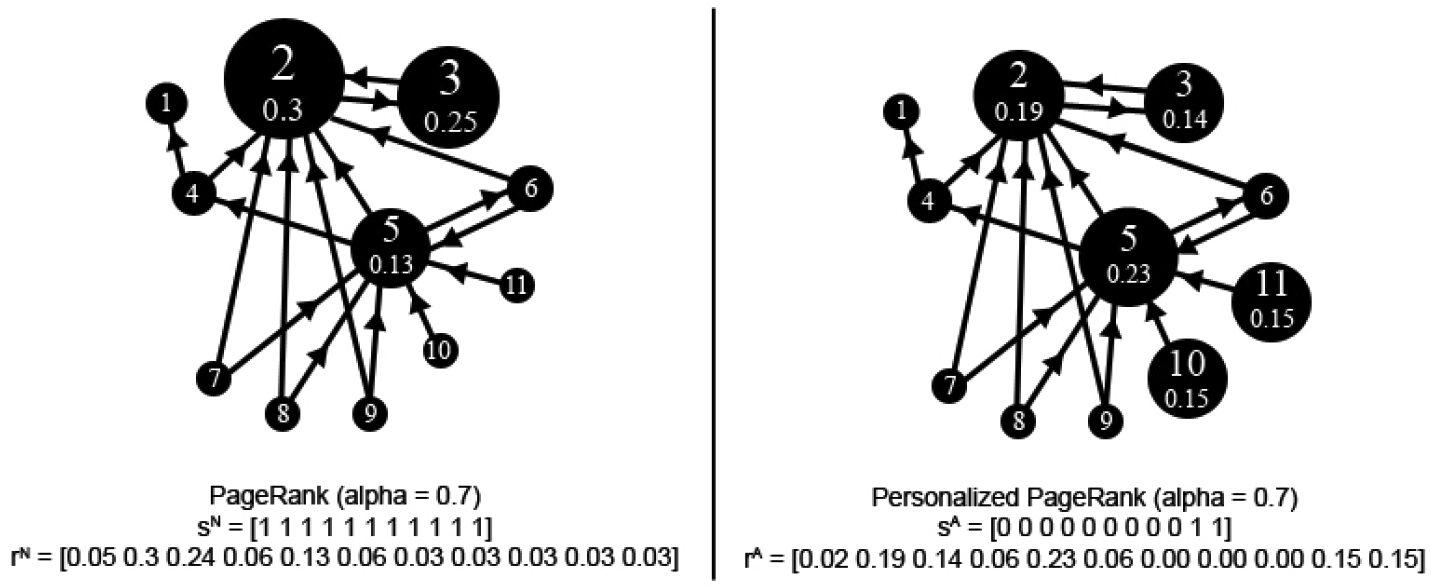
Illustrative example using differential sized vertices based on PageRank

The PageRank vector can be interpreted as the probability of landing at a given node by a random walker who jumps from vertex to vertex at each iteration. The random walker can with a certain probability *α* choose to teleport at random to any vertex in the graph instead of following an edge. The unit vector *ŝ* of the | V |- dimensional personalization vector *s* gives the probability of landing at a vertex if the walker teleports. *α* is known as the damping factor, and is a tunable parameter between 0 and 1.

The algorithm to compute the PageRank vector is described in Algorithm 1.

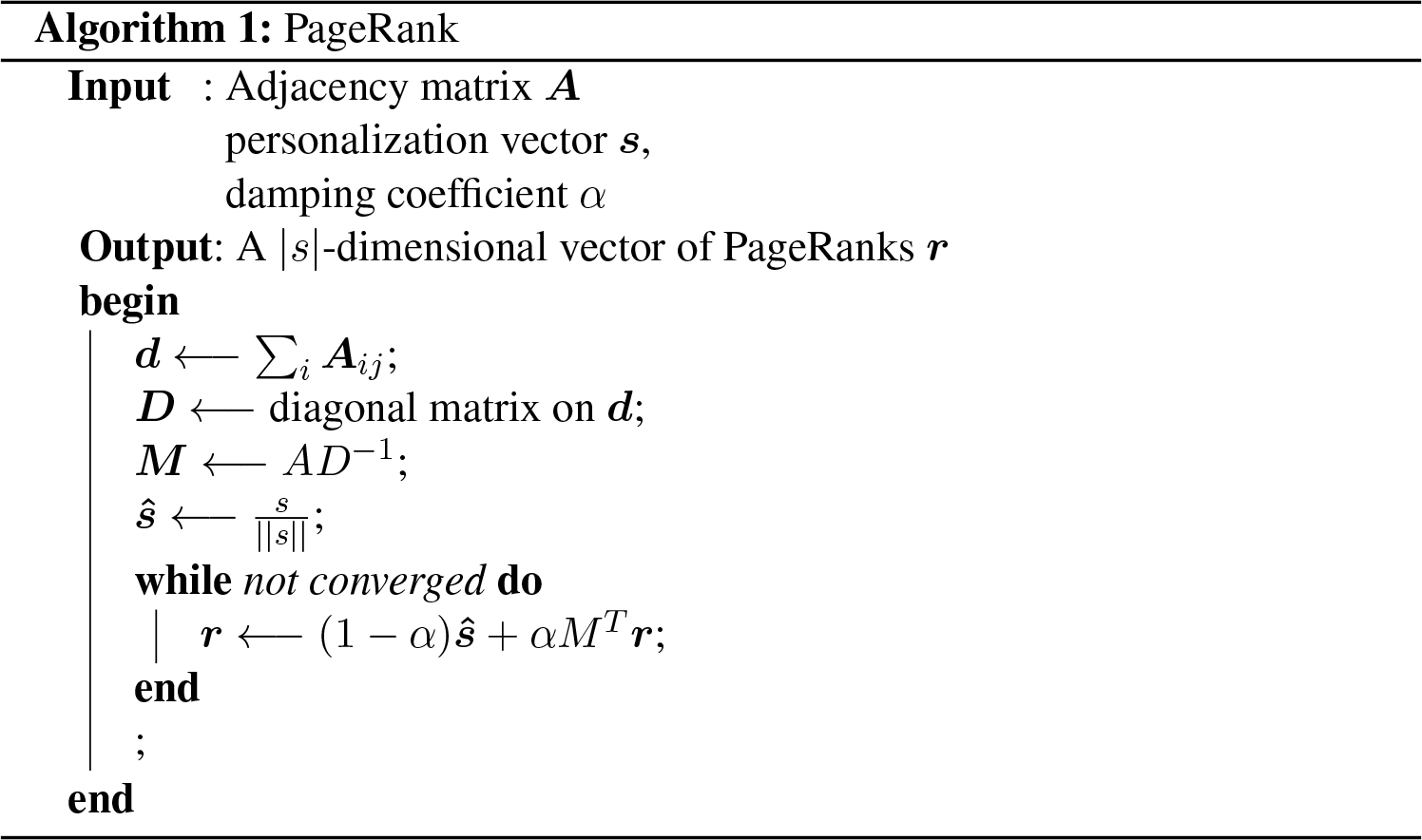

### 2.3 Pathway Analysis Model

Our model uses an available gene-gene interaction network. In this new approach, we treat gene scores such as fold changes or functional impacts as the extent to which the gene is disturbed and determines the personalization vector. Hence, each iteration of the page rank algorithm represents the propagation of disturbance through the gene interaction network. These are the components of our model

1. *G*(*V,E*) = The gene-gene interaction network, consisting of the set of interactions E between the set of genes *V*.
2. *p* = | V |-dimensional vector of gene p-values.
3. *f* = | V|-dimensional vector of gene fold changes.
4. *ϕ*(*p, f*) = a vector valued function that assigns weights to given vector of genes;
5. *α* = damping factor; 0 ≤ *α* ≤< 1
6. *r* = rank vector, computed using the PageRank algorithm using the quantities *G*(*V, E*), the personalization vector computed on the basis of *ϕ*(*p, f*) and *α* as input.

The interpretations of the components are as follows

- *ϕ* determines the personalization vector *s*, which indicates the bias of interaction of network towards various genes. The bias is directly proportional to *s*.
- *α* = damping factor serves dual purpose [11]. It can be interpreted as our faith in current graph structure and it can also be used to adjust the bias induced by personalization vector. A very small value (close to zero) indicates that we don’t trust the current graph structure and any gene would randomly interact with any other gene in the network irrespective of the edges in the graph. The magnitude of *α* also affects the rate of convergence of the algorithm. The rate of convergence is inversely proportional to the magnitude of *α*. *α* can also be used to model uncertainty and incompleteness of the gene interaction network. Most networks use value of around 0.85. We would recommend a values smaller than that (in range of 0.7 to 0.75) to compensate for the incompleteness of the interaction graph-structure.
- *r*= rank vector represents the final extent to which each gene is affected.

Thus, the page rank model takes into account the interconnected nature of the data and it can compensate for the incompleteness of the interaction graph structures.

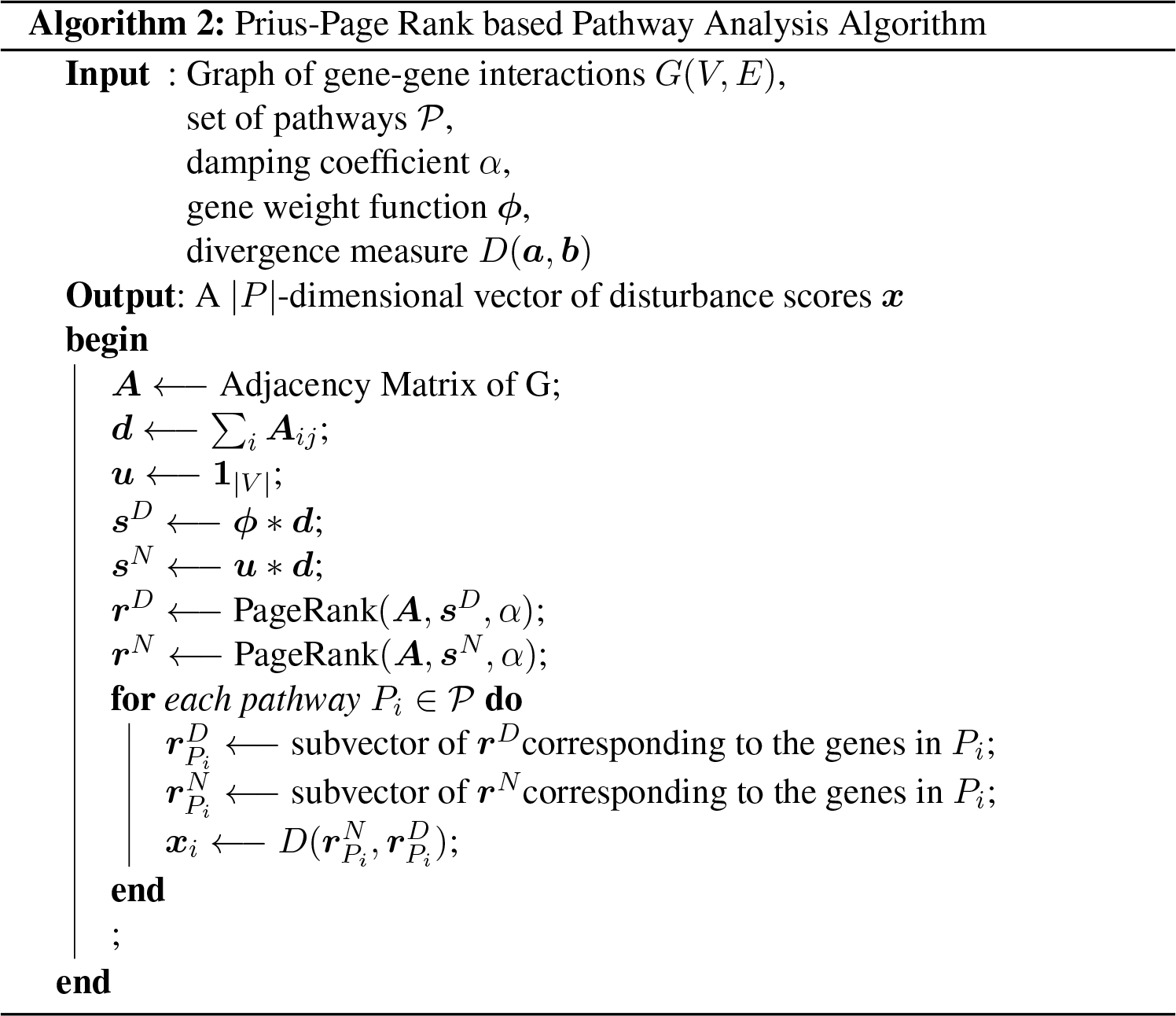

## 3 Prius - An Algorithm for Pathway Analysis

Here, we present an approach Prius for pathway analysis which differs from current enrichment based approaches. We would order the pathways based on how distribution ranks of the genes in a given pathway changes compared to itself and in overall context. For this purpose, we compute two different page ranks. These correspond to the following two scenarios

1. The term 1_|V|_ corresponds to a situation where the network has no bias.
2. The term *ϕ*(*p, f*) decides the extent to which each gene is disturbed and creates proportionate bias.
3. For both of the vectors, each entry is multiplied by the *outdegree* of the corresponding gene in the graph. The resulting vectors *s^N^* and *s^D^* are used as the personalization vectors for the normal and diseased condition respectively.

a. 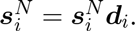.
b. 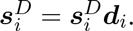.
c. This makes sure that the affected genes with higher outdegrees disturb the graph more. Without this correction, the disturbance caused by a gene with higher degree gets diluted.

Here, we use the same interaction matrix *M* and damping factor *α* in both these scenarios. Corresponding to the personalization vectors *s^N^* and *s^D^* we get two rank vectors *r^N^* and *r^D^* respectively. Now, we compute the distance of these two vectors for each pathway using an appropriate distance metric. The pathways are sorted in descending order of absolute distance to quantify relative affectedness of the pathways. This provides the appropriate prioritization of the affected pathways. We prefer not have any cutoff to remove pathways. In order to compute relative affectedness, we can compute divergence value of each pathway and sort pathways in descending order of pathways. A p-value can be computed from KL divergences if necessary. The entire procedure is summarised in Algorithm 2.

## 4 Illustrations for fine tuning of various components

Following are the components which one can fine tune

1. *ϕ*(*p, f*): following are some possible definitions

a. *ϕ* = |*f* | - This uses absolute value of fold change
b. *ϕ* = (1 — *p*) * |f | - where 1 denotes vector of all ones. This uses both p-value and fold change.
c. *ϕ* = 1 — *p* - Here we ignore fold change and use only p-values
d. *ϕ* = −*log*(*p*) - This formulation will introduce nonlinear exaggeration of p-values while deciding weights.
e. *ϕ* = −*log*(*p*) * |f | - This uses both p-value and fold change but nonlinearly exaggerates impact of p-value. In short, one can combine multiple linear and nonlinear combinations of p-values and fold changes to decide weights of genes in personalization vector.
2. Divergence - Consider a pathway *P* with k genes *g*1,*g*2, …,*gk*, and let 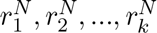 be their respective values in rank vector *r^N^* and 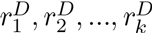 be their respective values in rank vector *r^D^*.

a. KL divergence - We can compute impact of disease on pathway *p* as Kullback-Leibler (KL) divergence of disease pathway vector *r^D^* from normal pathway vector *r^N^*. KL divergence is used to calculate distance between two probability distributions. In the given context, impact of disease on a pathway can be measured as

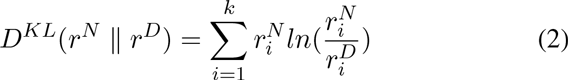

Where 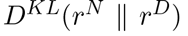 is divergence of *r^D^* from *r^N^*. KL divergence is a asymmetric measure. In certain scenarios, a symmetric measure might be desirable. Jensen-Shannon divergence can be used in these scenarios.
b. Mean Absolute Deviation (MAD): average of absolute fold changes of rank values of disease to normal.

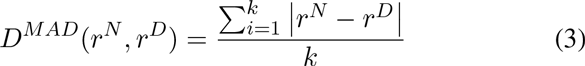
3. Interaction adjacency matrix M:

a. We have chosen to weigh all types of interaction equally. Different interactions (that is graph edges) can be assigned different weights depending on rate of interaction.
b. We have used unsigned matrix. However, one can model the interactions as signed edges. In such scenarios, variations of page rank from social network analysis based on trust and distrust propagation [12] can be used to compute rank.
c. M can represent heterogeneous graph that includes more cellular entities which can provide much more comprehensive model. In addition to genes, the following are some of the entities one can add

i. miRNAs can be added to the interaction network. Here, miRNAs can be connected to their targets. Edges could be signed.
ii. The promoters can be added and connected with corresponding gene. This can be useful to model impact of methylation.
d. M can even represent interactions among transcripts instead of genes. A gene can give rise to multiple transcripts and each transcript can code for different protein. With RNAseq, it is possible to estimate transcript description accurately. A transcript graph would provide finer level of information.

## 5 Materials and Methods

We use Reactome pathway database (Version 2015) to generate gene-gene interaction network. We use publicly available data from TCGA and GEO for our analysis. We demonstrate results of two analysis scenarios. The tool can be downloaded from https://github.com/bhatturam/prius

1. Personalized Analysis
2. Cohort Analysis

Damping factor is set at *α* = 0.7. For nodes that do not exist in the expression analysis, a default value of 1 was assumed. We use mean absolute deviation (Equation (3)) of the rank values to calculate pathway level disturbances.

### 5.1 Personalized Analysis

This analysis considers only a single patient’s data. Hence, a normal-tumor breast cancer sample pair (TCGA-BH-A0H7-11A-13R-A089-07, TCGA-BH-A0H7-01A-13R-A056-07) gene expression data is analyzed.

Here, *ϕ*(*p, f*) = | *f* |. Only the absolute fold change values are used for personalization. No p-value is calculated or used in this analysis. Top twenty pathways from this analysis are presented in Table 2.

**Table 2.** Personalized Analysis - Breast Cancer

| Reactome ID | Pathway |
| --- | --- |
| R-HSA-5638302 | Signaling by Overexpressed Wild-Type EGFR in Cancer |
| R-HSA-5638303 | Inhibition of Signaling by Overexpressed EGFR |
| R-HSA-1251932 | PLCG1 events in ERBB2 signaling |
| R-HSA-388479 | Vasopressin-like receptors |
| R-HSA-417973 | Adenosine P1 receptors |
| R-HSA-190370 | FGFR1b ligand binding and activation |
| R-HSA-1236382 | Constitutive Signaling by Ligand-Responsive EGFR Cancer Variants |
| R-HSA-1643713 | Signaling by EGFR in Cancer |
| R-HSA-5637815 | Signaling by Ligand-Responsive EGFR Variants in Cancer |
| R-HSA-5637810 | Constitutive Signaling by EGFRvIII |
| R-HSA-5637812 | Signaling by EGFRvIII in Cancer |
| R-HSA-2023837 | Signaling by FGFR2 amplification mutants |
| R-HSA-8853333 | Signaling by FGFR2 fusions |
| R-HSA-2033519 | Activated point mutants of FGFR2 |
| R-HSA-1253288 | Downregulation of ERBB4 signaling |
| R-HSA-2214320 | Anchoring fibril formation |
| R-HSA-444473 | Formyl peptide receptors bind formyl peptides and many other ligands |
| R-HSA-3000471 | Scavenging by Class B Receptors |
| R-HSA-8849469 | PTK6 Regulates RTKs and Their Effectors AKT1 and DOK1 |
| R-HSA-182971 | EGFR downregulation |

### 5.2 Cohort Analysis

This analysis is performed on small cell lung carcinoma samples. Microarray gene expression data from 60 pairs of normal and tumor is analyzed. This data is obtained from GEO( Dataset Record: GDS3837, Data Accession Series: GSE19804). p-value is obtained using a paired t-test.

Here, a personalization function based on both the computed p-values and the fold changes is used - *ϕ*(*p, f*) = *f* * (1 − *p*). Top twenty pathways from this analysis are presented in table 3

**Table 3.** Cohort Analysis - Lung Cancer

| Rank | Pathway |
| --- | --- |
| R-HSA-417973 | Adenosine P1 receptors |
| R-HSA-8851708 | Signaling by FGFR2 IIIa TM |
| R-HSA-442745 | Activation of CaMK IV |
| R-HSA-170660 | Adenylate cyclase activating pathway |
| R-HSA-167242 | Abortive elongation of HIV-1 transcript in the absence of Tat |
| R-HSA-72086 | mRNA Capping |
| R-HSA-6803529 | FGFR2 alternative splicing |
| R-HSA-167160 | RNA Pol II CTD phosphorylation and interaction with CE |
| R-HSA-77075 | RNA Pol II CTD phosphorylation and interaction with CE |
| R-HSA-170670 | Adenylate cyclase inhibitory pathway |
| R-HSA-997269 | Inhibition of adenylate cyclase pathway |
| R-HSA-1839126 | FGFR2 mutant receptor activation |
| R-HSA-111932 | CaMK IV-mediated phosphorylation of CREB |
| R-HSA-113418 | Formation of the Early Elongation Complex |
| R-HSA-167158 | Formation of the HIV-1 Early Elongation Complex |
| R-HSA-203927 | MicroRNA (miRNA) biogenesis |
| R-HSA-168325 | Viral Messenger RNA Synthesis |
| R-HSA-5655253 | Signaling by FGFR2 in disease |
| R-HSA-111957 | Cam-PDE 1 activation |
| R-HSA-444473 | Formyl peptide receptors bind formyl peptides and many other ligands |

## 6 Related Work

As explained in earlier sections, PageRank attempts to propagate an entity (belief, disturbance, perturbation-depending on the application) across the edges of the graph until steady state is achieved. The convergence is guaranteed if adjacency matrix is non negative. Certain variants model it for signed graphs, however due to lack of convergence they perform some maximum number of iterations and accept the final vector as rank vector. There are some examples of PageRank and its variants being used in NGS and pathway analysis in general. We briefly explain two such examples this section.

SPIA [13] combines both over representation analysis and perturbation analysis to come up with a p-value for a pathway. The perturbation factor for each gene is calculated as sum of signed log-fold change and the sum of perturbation factors of the genes directly upstream of the target gene, normalized by the number of downstream genes. So the perturbation propagation is similar to belief propagation. However this approach has two shortcomings. The perturbation propagation involves negative entries, so algorithm is not guaranteed to converge. Once the fold changes are captured in personalization vector, overrepresentations analysis becomes redundant.

DawnRank [14] attempts to rank mutated genes in a single cancer patient based on its potential to be a driver gene. Here, fold change serves as personalization component of PageRank. However, it uses *M* instead of *M^T^* in PageRank formulations. That means nodes with higher centrality values get ranked higher. Here, node centrality is interpreted as ability to affect other nodes in the graph.

## 7 Conclusion

In this paper, we proposed a novel and comprehensive approach, Prius for pathway analysis. This approach, enables all the genes to participate in the pathways analysis. The fold changes are treated as a disturbance received by gene and then PageRank perturbs the disturbance through the interaction network. The pathways are then ordered by their deviations from the normal state. This method thus overcomes the limitations posed by the current methodologies. Prius is also flexible as it offers users abililty to plug different algorithms of their choice for each stage of the work flow. Prius is a unique framework which offers a possibility of integrating heterogeneous entities like miRNAs and promoter into pathway analysis along with genes. We demonstrated the effectiveness of this method with two examples from cancer genomics.

## 8 Future Scope

In this paper, we proposed a new framework for pathway analysis which addresses some of the imminent issues of the state of the art. However, there is huge scope for improvement as follows.

- The current pathway databases are far from complete and have high degree of discordance. Though damping factor mitigates some of the uncertainty, a comprehensive pathway database will lead to more accurate analysis
- Rate of interaction is not currently available for all the interactions in the network. When rate of interactions will be available for all the interactions, Flux Balance Analysis (FBA) could be employed or page rank algorithm needs to be modified to accommodate rate of interaction.
- The interaction network can be enhanced by accommodating miRNAs, promoters and transcripts

